# funMotifs: Tissue-specific transcription factor motifs

**DOI:** 10.1101/683722

**Authors:** Husen M. Umer, Karolina Smolinska-Garbulowska, Nour-al-dain Marzouka, Zeeshan Khaliq, Claes Wadelius, Jan Komorowski

## Abstract

Transcription factors (TF) regulate gene expression by binding to specific sequences known as motifs. A bottleneck in our knowledge of gene regulation is the lack of functional characterization of TF motifs, which is mainly due to the large number of predicted TF motifs, and tissue specificity of TF binding. We built a framework to identify tissue-specific functional motifs (funMotifs) across the genome based on thousands of annotation tracks obtained from large-scale genomics projects including ENCODE, RoadMap Epigenomics and FANTOM. The annotations were weighted using a logistic regression model trained on regulatory elements obtained from massively parallel reporter assays. Overall, genome-wide predicted motifs of 519 TFs were characterized across fifteen tissue types. funMotifs summarizes the weighted annotations into a functional activity score for each of the predicted motifs. funMotifs enabled us to measure tissue specificity of different TFs and to identify candidate functional variants in TF motifs from the 1000 genomes project, the GTEx project, the GWAS catalogue, and in 2,515 cancer samples from the Pan-cancer analysis of whole genome sequences (PCAWG) cohort. To enable researchers annotate genomic variants or regions of interest, we have implemented a command-line pipeline and a web-based interface that can publicly be accessed on: http://bioinf.icm.uu.se/funmotifs.

## INTRODUCTION

Gene expression is orchestrated by regulatory elements that are sparsely distributed in the genome. The regulatory elements are activated or repressed by binding of TFs to specific DNA sequences known as motifs (1). Genetic variants associated with diseases and traits have been found systematically enriched in TF motifs (2). To prioritize the effect of noncoding variants, it is crucial to identify functional TF motifs. Veritably, only a fraction of the genome-wide detected motifs are predicted to be functional in any particular cell or tissue type.

Recent high-throughput sequencing technologies allow for detection of chromatin signals at a given time point (3). For instance, a single ChIP-seq experiment detects motifs that are bound by a specific TF (4). However, such experiment targets a single TF in a particular cell or tissue. Performing ChIP-seq experiments for all TFs in all cells and tissues to identify functional TF motifs is currently not feasible due to the lack of antibodies for many TFs and the time and cost issues for performing such experiments. In an approach to achieve this goal, thousands of ChIP-seq experiments in many cell and tissue types have been conducted in the ENCODE project (5, 6). Similarly, DNase1-seq experiments have been conducted to identify the accessible DNA regions in many cell and tissue types (7). Motifs located in the accessible regions are candidates to be bound by their matching TFs and thus they complement the information obtained from the ChIP-seq experiments. Overlaying chromatin signals that are detected to date in many large scale projects including ENCODE, the RoadMap Epigenomics and FANTOM provides a unique opportunity to identify functional motifs with the help of computational models (5, 8, 9).

Existing methods have proven to be useful for curating and reporting chromatin signals for given genomic regions (10–12). They have enabled researchers to prioritize genetic variants. For instance, RegulomeDB reports chromatin signals that overlap a list of input genomic regions or variants. The user is to determine whether signals from a relevant cell or tissue are present in the resulting list, however scanning large lists of annotations is a tedious task. Software tools such as CADD and AnnoVar have implemented methods to provide a summarized score for prioritizing genetic variants (11, 13). While they do provide scores for noncoding variants their main focus is on coding variants therefore the large sets of currently existing chromatin data remain unutilized.

To analyze the most crucial parts of regulatory elements, we have implemented a framework that utilizes large sets of chromatin data to annotate and score functionality of TF motifs.

## MATERIALS AND METHODS

### Motif prediction

We downloaded the curated non-redundant set of position frequency matrices (PFMSs) of 519 TFs (**Supplementary Table 1**) from the JASPR2016 CORE database (14). TF ChIP-seq peaks and DNase1 hypersensitive sites (DHSs) from all human datasets in the ENCODE data portal (n = 1,852, accessed on Dec 2016) were merged and converted to fasta format (15). Next, they were scanned for motif instances of the TF PFMs using FIMO (16). Motif instances that had a significant score (P-value < 1^e-4^) were obtained. Furthermore, to obtain high quality motif predictions, only the top significant motif instances (Z-score > 1) were retained from the significant motif instances of each TF.

### Annotation Collection

We curated high quality narrow peaks from TF ChIP-seq datasets in 18 cell lines (**Supplementary Table 2**). Peaks from multiple experiments were merged to generate a list of non-overlapping peaks per TF per cell line using Pybedtools (17). The same method was applied to curate DHSs in 19 cell-lines. The 15-state chromatin core models were downloaded for 17 cell types and tissues from The RoadMap Epigenomics data portal. Replication domains for 10 cell lines were obtained from Liu et al (18). We also downloaded the expression levels at promoters and enhancers that are based on CAGE peaks in the FANTOM project (19, 20). The median RPKM values for each TF were obtained from the GTEx project for the various tissue types (v6p, accessed on Mar 2017) (21). HiC data was obtained from Rao et al (22). We defined the boundaries of topologically associated domains (TADs) as contacting domains, and regions in between the boundaries were defined as loop regions. We provide scripts and manuals to generate the datasets described above. The scripts may also be applied to update the database content as new datasets become available, for instance, in ENCODE.

### Massively parallel reporter assays

We collected active regulatory elements that had been identified in massively parallel reporter assays (MPRA) (23–25). Elements that had a negative score were not considered. Motifs overlapping the elements were identified and for each element only the motif that had the largest activity score was kept. These motifs were used as functional motifs. To obtain non-functional motifs, we extracted motifs located in proximal promoters (1 Kb) of inactive protein-coding genes. Inactive genes were defined as genes that had TPM<1 in the ENCODE RNA-seq expression data in the respective cell lines. Finally, we annotated the functional and the non-functional motifs using data from matching cell lines. For instance, motifs that had a positive score from MPRA experiments in HepG2 cells were annotated with ChIP-seq peaks, DHSs, and other datasets from HepG2 and liver.

### Statistical modeling of functional annotations

The annotated sets of the functional motifs (n=10,793) and the non-functional motifs (n=11,198) were fed into a logistic regression model below (Equation 1). The input data to this model was in the form of a table where rows were motifs and the columns were the different annotations. The values in the table for each motif were preprocessed as follows. TF binding, DNase1 and CAGE were set to: “1” when there was an overlapping peak from a matching TF, an overlapping DHS and an overlapping CAGE peak, respectively, and “0” when the overlapping data was absent. TF expression level was log10 transformed. Dummy variables were introduced for the categorical values to incorporate replication domains and chromatin states into the model. Additionally, the 15 state chromatin states were summarized into 4 states by representing (**Supplementary Table 3**). The values of the dummy variables were set to 1 indicating an overlap with the motif and 0 indicating no overlap. In addition to the main TF, other TFs that co-localized at the same loci were counted as the number of other TFs.

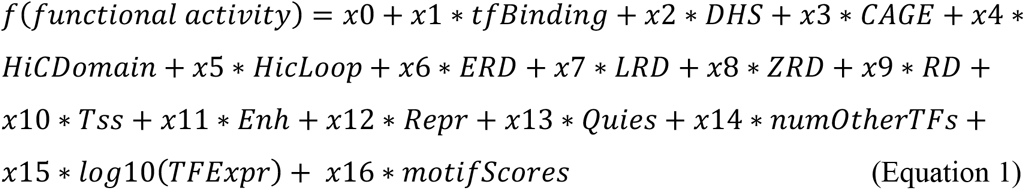

### Tissue-specific functional annotations

For each tissue, a set of relevant samples was defined (**Supplementary Table 4**). For each motif and in each tissue, annotations from the relevant samples were combined. For annotations with numeric values, the average value from the relevant samples was assigned; whereas for categorical values, the most common value among the relevant samples was assigned. Finally, in each tissue we imputed values for those annotations that lacked experimental datasets. The imputation was performed only for annotations that had experimental datasets in at least four tissues. For categorical annotations, the most common label was assigned to fill the missing value. Whereas for numeric features the mean value was taken from the tissues that had the annotations. This is particularly effective for TF binding and DHSs because they are set to exist even when binding evidence is found in only one tissue. For this reason we conditioned presence of DHSs to detect functional motifs as depicted below.

### Computation of motif functionality scores

To obtain a summarized functionality score for each motif, we used coefficients (*x1, x2,…x16*) that were positive and statistically significant (P-value<0.05). The coefficients indicate log odds ratio (logOR). Next, for each motif we multiplied the preprocessed values of its supporting annotations by their corresponding coefficients and finally summed the products. For instance, assume motif *M*1 had a direct TF binding, a DHS peak, two additional bound TFs and its chromatin state was *Enh*. The functionality score for *M*1 was then computed as the sum of: *x*1 * 1 + *x*2 * 1 + *x*11 * 1 + *x*14 * 2. The value for *number_other_TFs* was restricted to a maximum of three to limit its influence on the summarized score.

### Identification of functional motifs

Since DHSs mark accessible DNA, we conditioned presence of DHSs on all motifs. Motifs belonging to unexpressed TFs were considered as non-functional. Next, motifs were considered functional if they had binding evidence from matching TFs whereas those with evidence of no binding were considered as non-functional. For motifs where no TF ChIP-seq data was available, we required a significant functionality score. The significance was defined based on distribution of scores from functional motifs (matching TF-peak and DHS) in the myeloid tissue. The average score in the functional motifs was 3.725 and the standard deviation was 0.587.

### Calculation of variant effects on motif affinity

Firstly, the database was searched for identifying motifs that overlap a given variant, i.e. mutated motifs. Secondly, the position of the variant in the motif was identified by subtracting the variant start position from the motif start position, if the motif is on the positive DNA strand; for motifs on the negative strand, the motif end position is subtracted from the variant start position. Thirdly, entropy delta was computed as change in information content of the mutated motifs based on the nucleotide frequencies in the PFM model of the corresponding TFs. Frequency of the variants’ alternative allele was subtracted from frequency of the reference allele. A positive value indicates motif disruption, and a negative value indicates motif enhancement. In cases of multiple nucleotide variants or structure variants the entropy was set to 1.

### Identification of candidate variants

We downloaded human DNA variants detected in the 1000 Genomes project from the Ensemble Variation database (release 90: ftp://ftp.ensembl.org/pub/grch37/release-90/variation/vcf/homo_sapiens/1000GENOMES-phase_3.vcf.gz). They included Single Nucleotide Polymorphisms (SNPs) and short indels. Variants that had multiple alternative alleles were discarded. The funMotifs *Python* interface was used to identify the mutated functional motifs. Motifs that were functional in any of the tissues were considered. Variants that increased or decreased affinity of the overlapping functional motifs by at least 0.3 were retained.

Next, significant eQTL SNPs were downloaded from the GTEx data portal v7 (21). The SNPs from each GTEx tissue were annotated based on a corresponding tissue in the funMotifs database (**Supplementary Table 5**). These were: blood, brain, breast, colon, esophagus, liver, lung, pancreas, prostate, skin, stomach, and uterus. The GTEx tissues that had no matching annotated tissues were excluded. Variants that disrupted or enhanced information content of the functional motifs were retained.

Finally, SNPs were downloaded from the GWAS Catalogue (accessed on 11 Dec 2017) (26). The ENSEMBL REST API was used to curate SNPs that were in linkage disequilibrium (LD) of the GWAS SNPs (window size: 500Kb, r^2^>0.8). The coordinates were converted to hg19 using liftOver. The variants were annotated in tissues relevant to the phenotype (**Supplementary Table 6**). Variants disrupting or enhancing functional motifs were retained from each tissue and for each phenotype.

To compare existing functional annotation methods, we used the set of eQTL SNPs from the GTEx data portal v7 (21). The SNPs were annotated using RegulomeDB, HaploReg and funMotifs. The haploR package was used to annotations from RegulomeDB and HaploReg (10, 12)(27). To identify candidate variants, we selected SNPs that overlapped a TF motif with a matching TF binding event as well as a DNase1 peak in the corresponding tissue. Additionally, we considered variants if the mutation changed the motif entropy significantly. For RegulomeDB, we considered variants with the score 1a, 1b, 2a and 2c in order to keep variants with the aforementioned evidences. However, RegulomeDB did not provide any numerical measurement determining the effect of the mutation on motif affinity. Candidate variants from HaploReg were further filtered to only keep those with a minimum of 30% log-odds (LOD) difference between the refence and the mutated motif. All candidate functional variants were considered in a tissue-specific manner (**Supplementary Table 5**) by utilizing data from ENCODE (http://genome.ucsc.edu/ENCODE/cellTypes.html#TOP) (15). For funMotifs we used thresholds introduced in this paper.

### Implementation of funMotifs

A PostgreSQL database was created to store the generated data. To increase the performance of database queries, the motifs were split into separate tables based on their chromosomes, and they were indexed based on their genomic coordinates using the integer-range method. A table was created per tissue to store the annotations and the scores of the motifs in that tissue. The framework was implemented in Python 2.7 and the code repository is made publicly available. A programming interface is provided to access the database content to analyze large sets of genomic variants. Also, an easy-to-use web interface is provided to analyze smaller data sets.

## RESULTS

### Motif annotation framework

We created a framework (funMotifs) to curate and integrate chromatin signals across cell lines and tissues from pre-defined data sources. funMotifs generates tissues-specific annotations by integrating signals from the set of samples assigned to each tissue. In cases where no data is available for a tissue, it imputes the average value from the other tissues. Next, it overlays the annotations onto a set of given TF motifs (**Figure 1**). A functionality score for each motif is computed based on the importance of the annotation features of the motif. The importance of the annotation features may also be separately provided to the framework. Finally, to facilitate usage of the annotated motifs, the framework generates a PostgreSQL database to store the functionality scores and annotations of the motifs per tissue.

**Figure 1:**
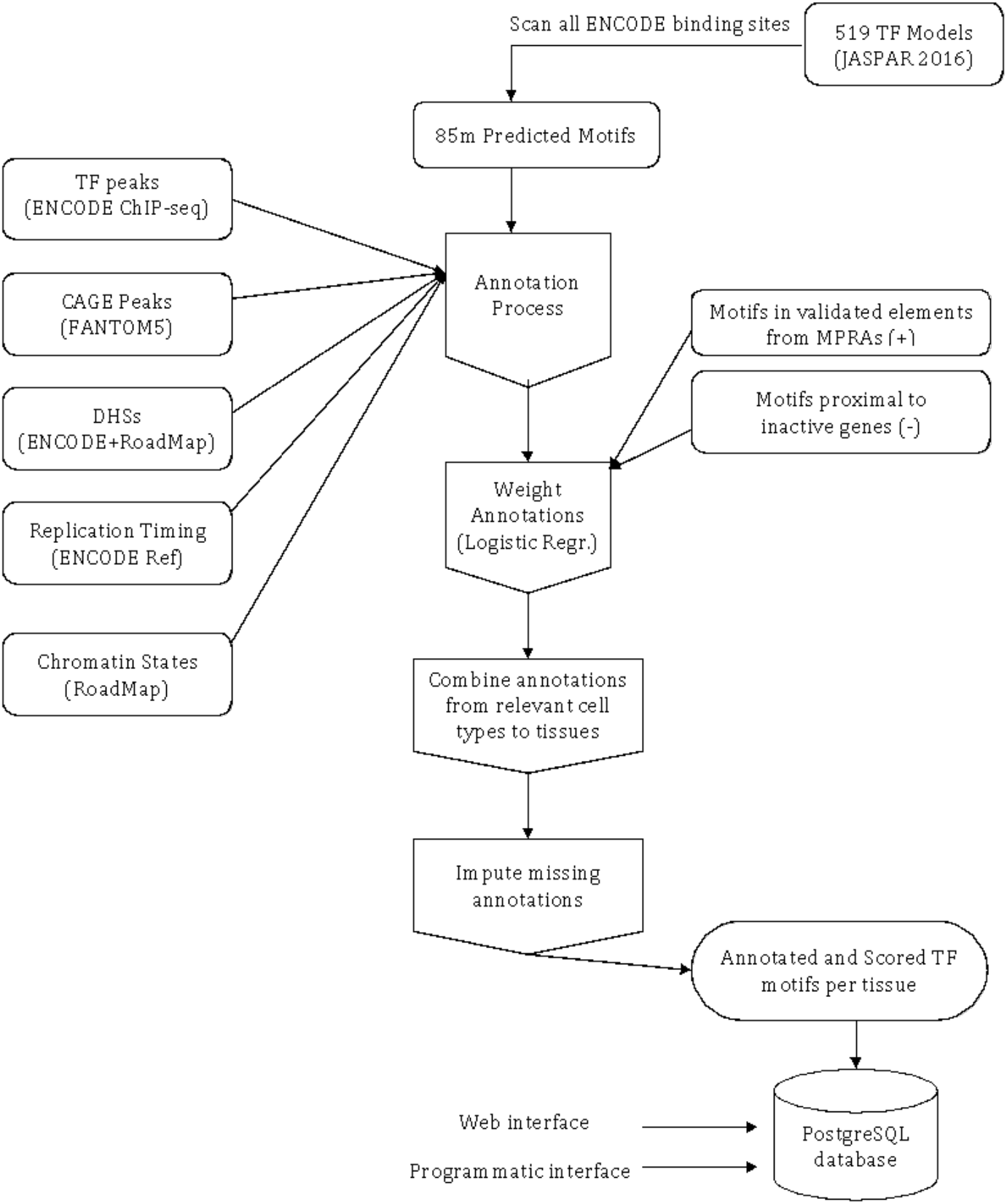
funMotifs framework pipeline to annotate motifs. Sequence motifs were predicted based on PFM models from the JASPAR database. Annotation datasets were collected from the listed data types. The resources used for each data type are indicated in the parenthesis. The annotations are overlaid to the motifs. Motifs in validated regulatory elements were combined with motifs near inactive genes. The positive and negative cases were used to build a logistic regression model for identifying functional motifs. The annotated and scored motifs are stored in a database. Programmatic and web interfaces are provided to utilize the annotated motifs.

We applied the funMotifs framework, to annotate sequence motifs (n = 85,459,976) of 519 TFs (**Supplementary Table 1**) predicted in the entire potential regulatory landscape of 105 cell lines. Overall, the motifs covered 14% of the human genome. Subsequently, to annotate the motifs across 15 tissues we curated annotations for the samples listed in (Materials and Methods, **Supplementary Table 2**). The annotations included high quality TF peaks (n = 14,902,767) obtained from 1,604 ChIP-seq datasets in 18 cell lines. The curation was restricted to narrow peaks, and when available, we used peaks that had been identified through the Irreproducibility Discovery Rate method to ensure high data quality (28). We also obtained DHSs (n=6,676,179) from 21 cell lines, chromatin states, HiC domains and loops, replication domains, TF expression, and CAGE peaks (See Methods and Materials).

Next, we identified the importance of each annotation feature by training a logistic regression model on validated regulatory elements identified in MPRA experiments (Methods and Materials). As expected, presence of a motif matching TF peak provided evidence for functionality of the motif (log-odds ratio=1.4) (**Figure 1**). Additionally, motifs located in active elements such as enhancers and transcription start site (TSS) regions also had a higher odds ratio to be active (logOR ~ 0.91). Whereas quiescent regions had a negative enrichment at logOR of −0.3. Interestingly, early replicated regions also had a higher enrichment of active motifs (logOR = 0.79) compared to late replicated regions (logOR = 0.01). CAGE peaks (logOR = 0.2) and binding of other TFs referred to extra indications for functionality of the motif (logOR = 0.14). Functionality scores for the motifs were computed in each tissue by using the positive coefficients from the logistic model (Methods and Materials).

### Motifs across the genome

The functionality scores ranged between 0 and 4.31 where low scores indicate no functionality and high scores indicate functionality of the motif. As expected, majority of the motifs had very low functionality scores in all the tissues (**Figure 2a**). The scores varied largely between the motifs of different TFs. For instance FOXA1 and MAFK had a large number of motifs with ‘outlier’ scores whereas CEBPB and CTCF had higher scores for motifs in their fourth quartiles across the tissues (**Figure 2b**). To select functional motifs we conditioned on presence of DHS, and a score larger than 2.55 or a matching TF peak. The threshold was based on score distribution of motifs that had matching TF peaks and DHSs in the myeloid tissue since it had the largest size of annotation data (**Figure 2c**).

**Figure 2:**
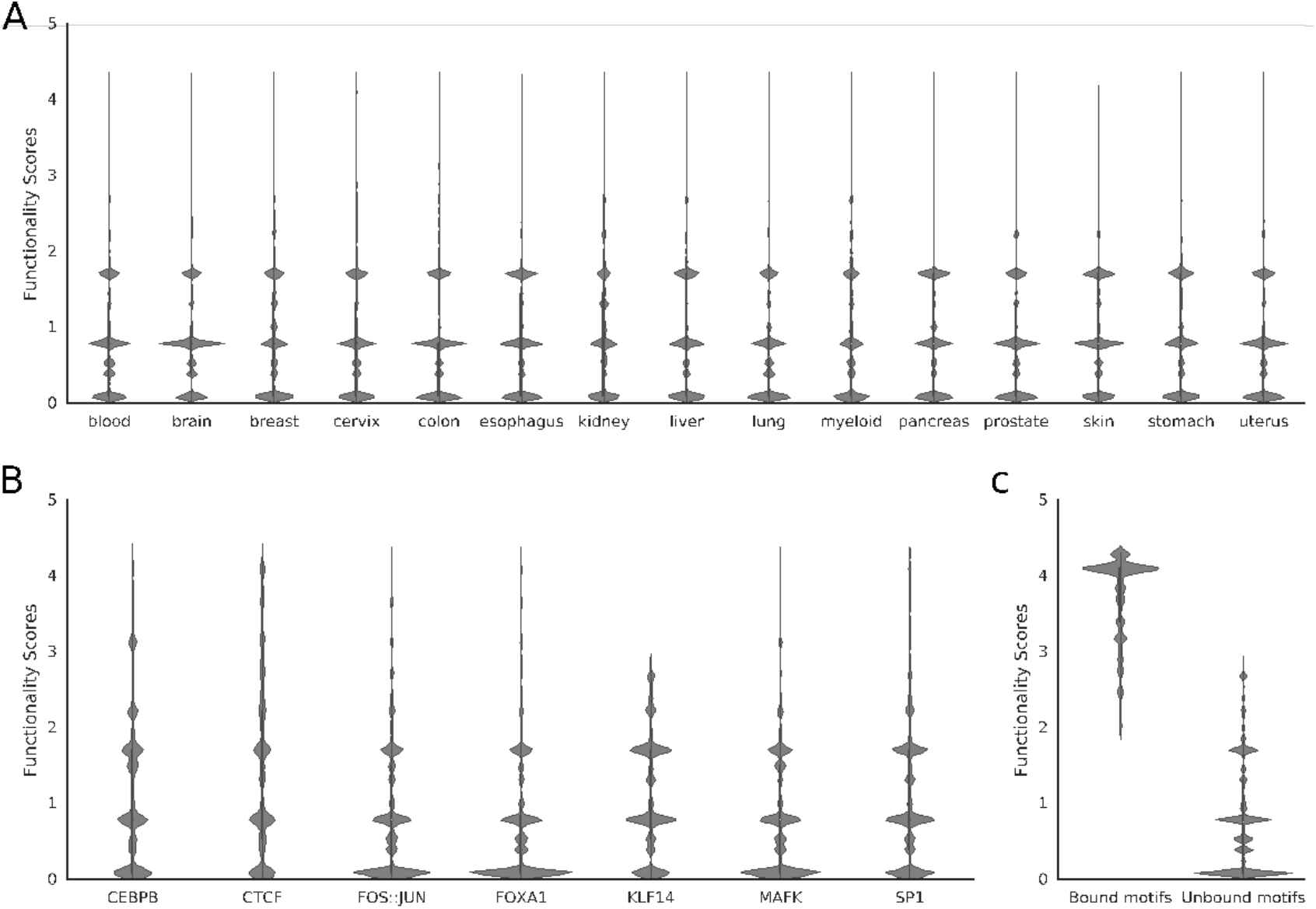
Distribution of functionality scores across tissues and TFs. (A) Distribution of functionality scores per tissue. Scores from motifs of all TFs are combined. Majority of motifs have scores below 2 which indicates a nonfunctional score. (B) Functionality scores in motifs of selected TFs. The scores across the tissues are combined. (C) Functionality scores in motifs that are bound by matching TF peaks and DHSs in myeloid tissue and those that are not bound.

Overall, less than 1% of the motifs were found functional across all tissues. Altogether, they covered 0.74% of the genome and formed 1,660,964 functional sites across the tissues. Merging functional motifs within 300bp from all the tissues indicated 364,466 regulatory elements covering 3.1% of the genome. All TFs had a small fraction of their motifs functional and there was no positive correlation between the number of sequence motifs and the number of functional motifs (**Figure 3a**). FOXP1 and NFATC2 each had more than 1 million sequence motifs, but only around 30k of them were predicted to be functional. In contrast, KLF5 had about 427k sequence motifs but it had the highest number of functional motifs in the genome (n=68,695). Based on our model, presence of TF binding was a strong indicator for functionality of the motifs. However, in most cases the number of functional sites was slightly smaller than the number of motifs that overlapped TF peaks mainly due to conditioning on the presence of DHSs (**Supplementary Figure 1 a-c**). Furthermore, our method could also identify functional motifs for TFs where no ChIP-seq data was available, as exemplified by KLF14 in the blood tissue (**Supplementary Figure 1a**).

**Figure 3:**
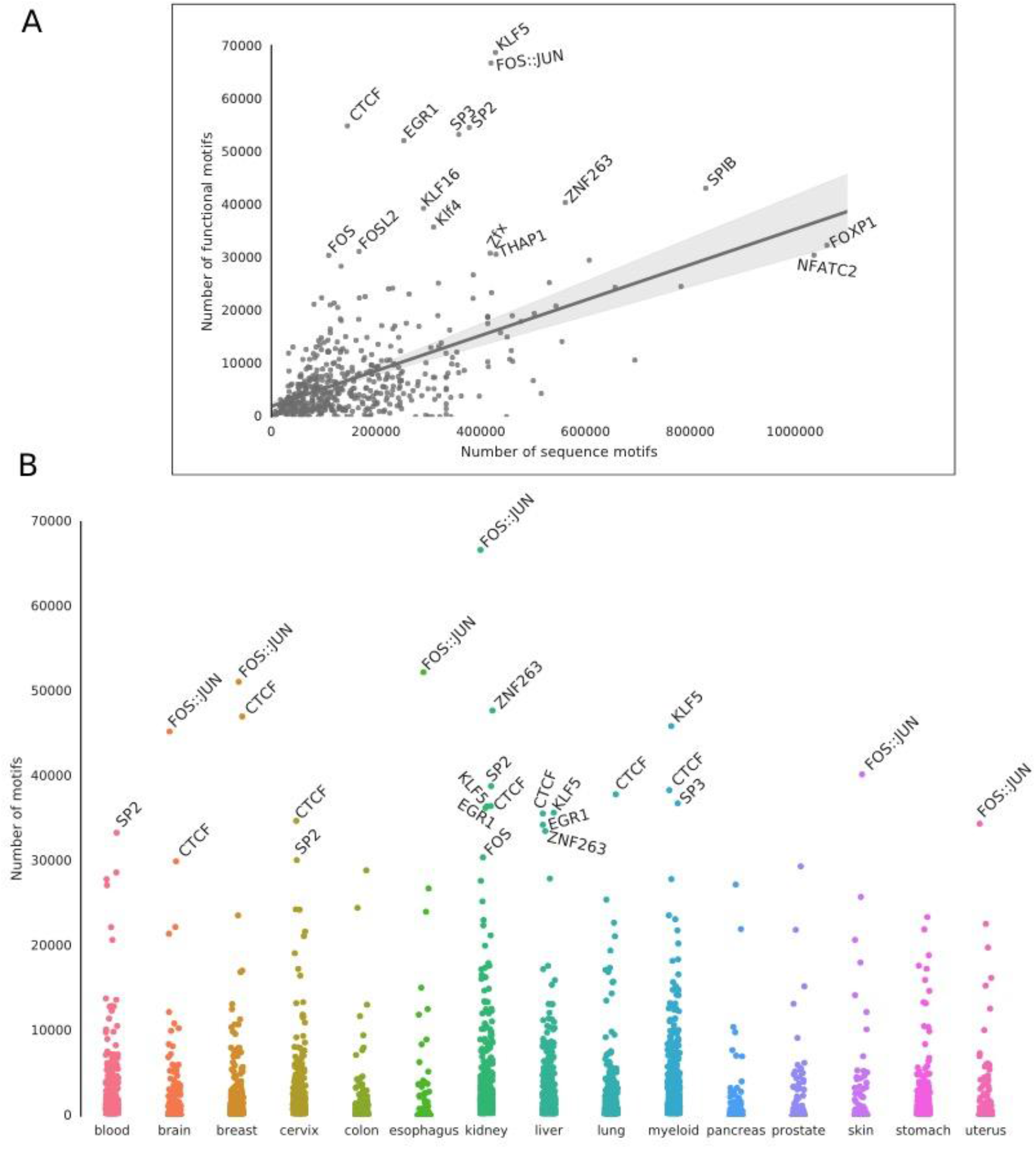
Frequency of functional motifs per TF. Motifs of each TF that are identified as functional are merged across the tissues. Each dot shows the number of motifs predicted from the genome-wide scan (x-axis) and the number of functional sites for a TF (y-axis). (B) Each column represents a tissue and each dot represents the number of functional motifs per TF in the corresponding tissue. TFs that had more than 30,000 functional motifs were labeled.

There were large differences between the number of functional motifs among the TFs. Promoter binding factor AP-1 (FOS::JUN) that regulates expression of many genes had the largest number of functional motifs after KLF5. These were followed by CTCF (n=54,834), which is a multifunctional TF and it is highly expressed in all tissues (**Figure 3a**) (29). 249 TFs had more than 5,000 functional sites and 113 of them had more than 10,000.

### Tissue-specificity of TF motifs

The number of functional motifs in each tissue was highly dependent on the number of input assays. For instance, there were 789,191 functional sites in myeloid tissue that had the largest number of TF peaks as input. This was also the case for kidney (n=763,141 sites), liver (545,236), and blood (411,649) that had a high number of ChIP-seq datasets. In contrast, pancreas had the least number of functional sites (n=98,836). This indicates that further tissue specific functional annotations need to be provided to identify all functional sites.

The functional sites were highly tissue-specific. Interestingly, 52.6% of them were found in only one tissue and 16% in only two tissues. However, since the shared sites were mostly between the tissues that had the largest annotation datasets, namely myeloid, kidney, liver and blood, we expect a higher number of sites shared between the tissues. Overall, 1.5% of the sites were shared among all 15 tissues.

AP-1 and CTCF motifs were the most functional in the majority of the tissues (**Figure 3b**). In contrast KLF5 motifs were highly frequent only in myeloid, liver and kidney. SP2 motifs were also found highly functional in blood, cervix, and kidney.

Additionally, we looked for enrichment of motifs in various chromatin states. Since our model was giving higher scores to motifs in TSSs and enhancers, they had the highest enrichment for functional motifs. Overall, 36% of the functional motifs were located in TssA, and 22% were located in enhancers whereas only 9% were located in quiescent regions. However, this enrichment varied between motifs of different TFs. For instance in liver, the promoter binding protein families (EGR, KLF and SP) were all highly enriched in TSS-like states (**Figure 4**) whereas CTCF was highly enriched in quiescent and repressed states. The same enrichment was also observed in other tissues including myeloid and blood tissues (**Supplementary Figure 2 and 3**).

**Figure 4:**
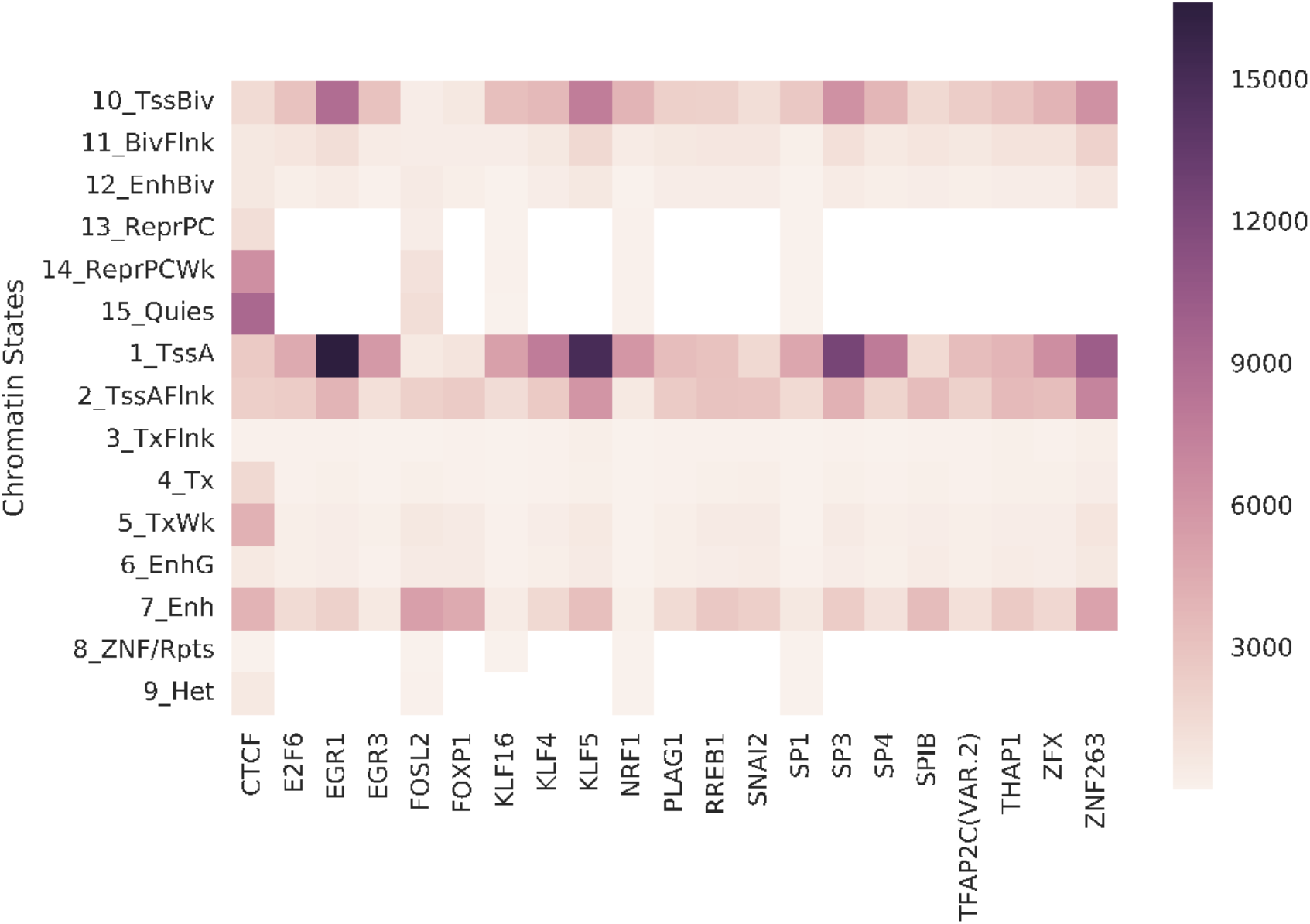
Enrichment of functional motifs in chromatin states in liver tissue. The cells show the number of functional sites per TF per chromatin state.

### Prioritization of noncoding variants

Variants detected in whole genome sequencing and genotyping studies are vast and sparse in the noncoding genome. However, distinguishing variants that affect gene regulation is a difficult task due to the large search space. Generation of the annotated motifs allowed for a systematic prioritization of noncoding variants. We built a web-based interface to enable researchers upload a list of variants and identify affected motifs. The web-based system uses the database that contains annotations and scores of the motifs in all 15 tissues. To get tissue specific annotations for the affected motifs, the user may select a tissue that is most relevant to the disease or condition in question. Alternatively, a *Python*-based interface is also provided to allow for analysis of large sets of variants. For each input variant, the program returns the list of motifs that overlap the variant. The variant is then given a final score to indicate its deleteriousness. The score is computed as functionality of the overlapping motif, and the variants effect on disrupting or enhancing affinity of the motif. The user is then able to choose candidate variants from the resulting list of scored variants. The annotation features allow for further in-depth analysis on the role of noncoding variants in the genome. For instance, enrichment of noncoding variants in the various chromatin states, replication domains and HiC contacting domains are all provided from the database.

We applied the funMotifs framework to analyze somatic mutations from 2,515 cancer whole genome sequences in the PCAWG project (30). By limiting the set of 25,277,342 mutations to those that had potentials to affect functional motifs in tissues corresponding the cancer types, we were able to obtain a prioritized set of mutations containing less than 1% of the initial set (Umer et al. *Manuscript)* (31).

In comparison to the existing methods, funMotifs provides larger and more specific annotations for TF motifs. In **Table 1**, we compare the main features of funMotifs to RegulomeDB, HaploReg and AnnoVar (10–12). funMotifs differs from the currently available databases in two ways. Firstly, it uses tissue-specific annotations to score the effect of variants, and secondly, it is specifically designed to analyze mutations in TF motifs. RegulomeDB provides scores based on the annotations but unlike funMotifs, the scoring is not mathematically computed. Rather it is a discrete categorization of the annotations. AnnoVar combines scores across several methods with functional annotations to prioritize variants. However, its ability for analyzing variants in TF motifs is limited because the user has to provide ChIP-seq peaks. Similarly to funMotifs, CADD integrates multiple annotations to generate a deleteriousness scores for genomic variants (13). However, its annotation sets and features are limited and many of the features are to predict the effects of coding variants. Overall, none of the existing tools provides resources to analyze the effect of genomic variants on motifs in consideration of binding of matching TFs. Also, information from CAGE peaks, replication timing, and HiC data have not been utilized by the existing methods.

**Table 1:** Comparison to existing annotation databases

| Features | funMotifs | RegulomeDB | HaploReg | AnnoVar | CADD |
| --- | --- | --- | --- | --- | --- |
| Genome wide | No | Yes | Yes | Yes | Yes |
| Functionality score | Yes | Yes | No | Yes | Yes |
| Functional annotations | Yes | Yes | Yes | No | No |
| Tissue Specificity | Yes | No | No | No | No |

We evaluated the ability of funMotifs for detecting functional variants in comparison with HaploReg and RegulomeDB. The GTex SNPs were annotated by each tool in a tissue specific manner (Methods and Materials). The highest number of candidate variants were detected by funMotifs (5,375), where HaploReg and RegulomeDB detected 1,972 and 1,503 variants, respectively (**Supplementary Tables 7-9**). Only 84 candidate SNPs were detected by all databases. The overwhelming majority of the candidate variants were detected individually by each method (**Supplementary Figure 4**). RegulomeDB detected candidate variant in eight tissues (blood, brain, breast, colon, liver, lung, pancreas and skin), HaploReg in nine tissues and funMotifs in twelve tissues. The funMotifs approach achieved the highest number of the candidate functional SNPs in all tissues. The distribution of the variants between tissues was similar between the two remaining tools (**Supplementary Table 10**). Notably, only cell lines from Tier 1 and Tier 2 from the ENCODE project were used to annotated the motifs in funMotifs whereas cell lines from Tier 3 were also used to annotate variants in HaploReg and RegulomeDB (Methods and Materials, **Supplementary Table 4**).

### Enrichment of genetic variants

Next, we sought to identify the effect of variants on TF motifs. We curated 84,346,970 SNPs and short indels from the 1000 Genomes project in 2,504 samples (32). In total 12,677,702 variants overlapped TF motifs (Methods and Materials). We identified 463,758 functional motifs overlapping 350,646 candidate functional variants with disruptive effects (0.42% of all the variants) (**Data File 1**). 26.5% of the variants were common with a minor allele frequency (MAF) > 0.01 and the remaining 73,5% were rare with a minor allele frequency lower than 0.01 in all the populations. 43.3% were C>T or G>A variants indicating large effects on methylation sites. Notably, nearly 7% of the variants were found in position 14 and 15 of the CTCF motif. Mutations at this locus have been attributed to impair of the DNA excision repair machinery (33). Interestingly, highly conserved positions in motifs of SP1, KLF5, CTCF, and EGR1 were the next most mutated. SP1, KLF5, and EGR1 bind to promoter regions and regulate the expression of a large number of genes that are involved in variety of essential cell processes. Since majority of the variants affected methylation sites of the motifs they may play a role in epigenetic regulation. Additionally, 151,203 variants were found increasing information content of the functional motifs indicating enhancement of TF binding, and 23,5% of them were common variants. Overall, 24,5% of the variants resided in enhancer elements, and 93% were in early replicated domains.

We next evaluated the effect of eQTL SNPs on TF motifs. Significant eQTL SNPs (n=2,219,330) were downloaded from the GTEx project for the annotated tissues (Materials and Methods). In total 5,375 candidate SNPs were identified in the analyzed tissues (**Supplementary Table 7**). The largest number of candidates was identified in blood (n=2,473), followed by lung, brain, and breast. The identified SNPs were in significant associations with 5,383 genes. Notably, up to 59 candidates were associated with HLA genes. In contrast, 59.4% of the genes were associated to only one candidate SNP. Interestingly, 63% of the eQTL SNPs were in TSS chromatin states, and 253 SNPs were in TSS states in some tissues and in enhancer states in others.

GWASs have reported thousands of SNPs to be associated with various diseases and traits. However, many of them remain uncharacterized due to the complexity of the noncoding genome. In order to identify potential candidates, we curated GWAS SNPs associated to several phenotypes and terms, and for each term we assigned a matching tissue (**Supplementary Table 6**). Other SNPs that were in LD with the GWAS SNPs (r^2^>0.8) were also included. Overall, we identified 1,083 candidate SNPs affecting function TF motifs (**Supplementary Table 11**). Phenotypes that were linked to myeloid (n=499 SNPs), blood (n=414) and liver (n=240) tissues had the highest number of candidates (**Supplementary Table 12**). 76% of the candidates were common variants (MAF>0.01 in the 1000 genomes project). Interestingly, 265 candidates were also among the eQTL SNP candidates. The identified candidates are expected to have impacts on the observed phenotypes through differential gene regulation.

Examples of candidate GWAS SNPs that had additional evidence from the literature included SNP rs8103622 that was identified as an important candidate functional variant in four tissues: blood, breast, liver and myeloid (**Supplementary Table 11**). rs8103622 was in strong LD with four SNPs from the GWAS catalog, two of them (rs4808801 and rs7258465) were shown to be breast cancer risk variants (34). The eQTL analysis for the breast tissue indicated association with a SSBP4 gene, which is a tumor-suppressor gene (35). The candidate SNP changed binding affinity of a CTCF motif (**Figure 5**). The SNP resulted in a significant allelic imbalance between alleles C and T, nearly 79% and 21%, respectively in a set of MC7 cell lines (**Supplementary Figure 5 and Supplementary Table 13**). Our findings corresponded to the results for rs8103622 in (36). Moreover, we checked the presence of DHSs and CTCF signals for the remaining tissue types: blood, liver and myeloid. In all cell lines DNase1 peaks were shown, however only for breast cancer and blood cell lines we observed a CTCF occupancy on a noticeable level (**Supplementary Figure 6**). In addition, two remaining GWAS SNPs: rs4808136 and rs28375303 in LD with rs8103622 were associated with blood metabolite levels and a monocyte count (37, 38). Above findings demonstrate that the candidate allele-specific SNP rs8103622 is a potentially functional variant in breast cancer and blood. Additionally, rs9271562 was identified as a candidate functional variant affecting the CTCF motif in blood with a functional score of 4.1. Candidate SNP rs9271562 was in LD with the GWAS SNP rs3129763 that has been associated to multiple sclerosis (**Supplementary Table 11**). Interestingly the eQTL analysis associated the SNP with four genes from the HLA family, one of them: HLA-DRB1 that plays role in multiple sclerosis (39, 40). A high distributive effect (entropy value 0.96) and presence of TF binding at the loci may influence gene deregulation and risk of multiple sclerosis (**Supplementary Figure 7a**). Finally, we investigated the candidate variant rs226198 in the breast tissue (**Supplementary Table 11**). rs226198 has been associated with two breast cancer SNPs from the GWAS catalog: rs146817970 and rs7707921 (34). The variant overlapped a MYC:MAX heterodimer motif, the MYC:MAC dimer is crucial for binding of MYC to the TF binding site (41). Both MYC and MYC:MAX are known as oncogenes in many cancer types, including breast cancer (42–44). Genes: ATP6AP1L, ATG10 and RPS23 were found affected by the candidate SNPs. Moreover, the candidate SNP was located in a chromatin accessibility region (**Supplementary Figure 7b**). Results presented here are promising, however, further studies need to be conducted to evaluate their impact on the genes *in vivo*.

**Figure 5:**
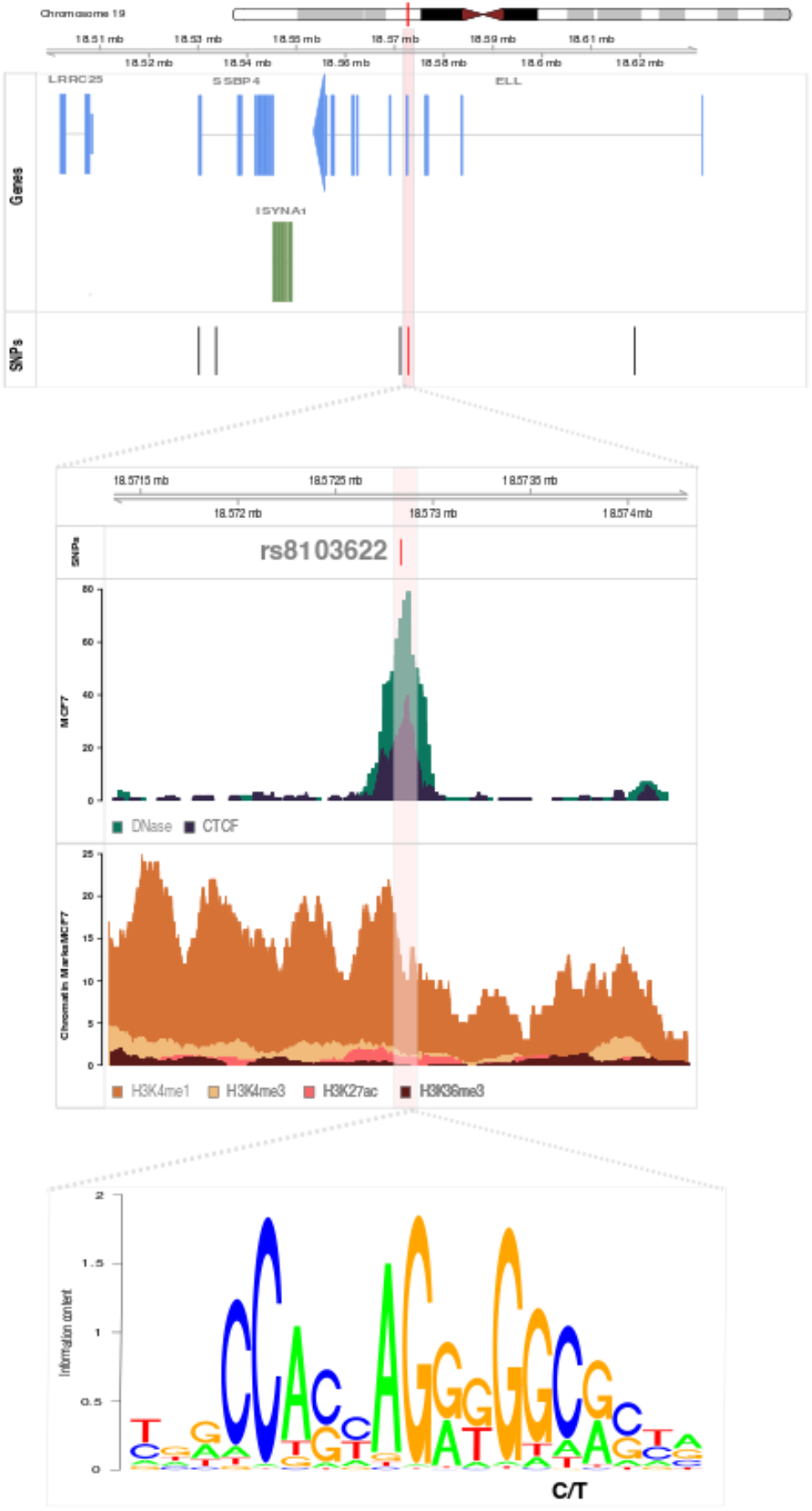
Overview of the candidate functional GWAS SNP rs8103622 in breast tissue. Genes box: blue genes are genes associated with the candidate SNP using eQTL SNPs. Green gene is not associated with the candidate SNP according to eQTL SNPs. SNPs box: red line indicates position of the candidate SNP rs8103622, black lines are GWAS SNPs in LD with the candidate SNP.

## DISCUSSION

Noncoding variants deregulate gene expression by altering regulatory regions. The functionality of regulatory elements is determined by the DNA motif sequence. Identification of functional DNA motifs is essential to annotate gene regulation. funMotifs framework was used to combine large sets of annotation features obtained from ChIP-seq, DNase1-seq and other experiments. A logistic regression model was applied to score motifs of more than 519 TFs across the genome.

The resulting data showed that less than 1% of the genome is covered by TF motifs with indications that more remain to be discovered since the analyses are based only on a subset of all human TFs (45). Also, we expect that the number of functional motifs for some of the TFs be eventually larger than reported due to the presently limited ChIP-seq datasets in many cell and tissue types. Furthermore, conditioning on DNase1 sites excluded motifs that might be functional outside the open chromatin (46). The sets of functional motifs showed high degrees of tissue specificity as 52% of the identified motifs were unique to a single tissue. This finding emphasized the importance of characterizing noncoding variants based on annotations from cell and tissue types that are relevant for the phenotype.

Furthermore, to enable researchers identify motifs in genomic regions of interest, we have built an online interface. The web interface provides access to annotations in fifteen tissue types. They provide means to prioritize noncoding variants regarding their effect on TF motif affinity. Applications of the tool identified potentially functional variants in cancer and normal genomes. Overall, we identified 350,646 candidate variants from the 1000 Genomes project, 5,375-candidate eQTL SNPs from the GTEx project, and 1,083 candidates from the GWAS catalogue. The enriched TF motifs provide insights into the regulatory role of genetic variants. Majority of the identified candidates are expected to have impacts on differential gene regulation. In the study we presented examples of such candidate SNPs, which overlapped potentially functional motifs in different tissues. However, further studies need to be conducted to evaluate their impact on the genes *in vivo*.

While we have provided tools and resources to analyze the effect of variants on TF motifs, variants that lead to creation of de-novo motifs also need to be taken into account. Altogether, the functional activity annotations of the motifs provide further insights into our understanding of the gene regulation process in the human genome.

## FUNDING

This work was funded by Uppsala university [H.M.U]; institute of Computer Science, Polish Academy of Sciences and SYMFONIA project from Polish National Science Centre [J.K.]; grants from the Swedish cancer foundation (No. 160518) [C.W.].

## Conflict of interest statement

None declared.

## Supplementary Material

### Supplementary Tables are available on

https://figshare.com/s/e6d3501d5a36617893b5

### Data File 1 is available on

https://figshare.com/s/073c985fd131f78fb9c5

### Supplementary Figures

**Supplementary Figure 1:**
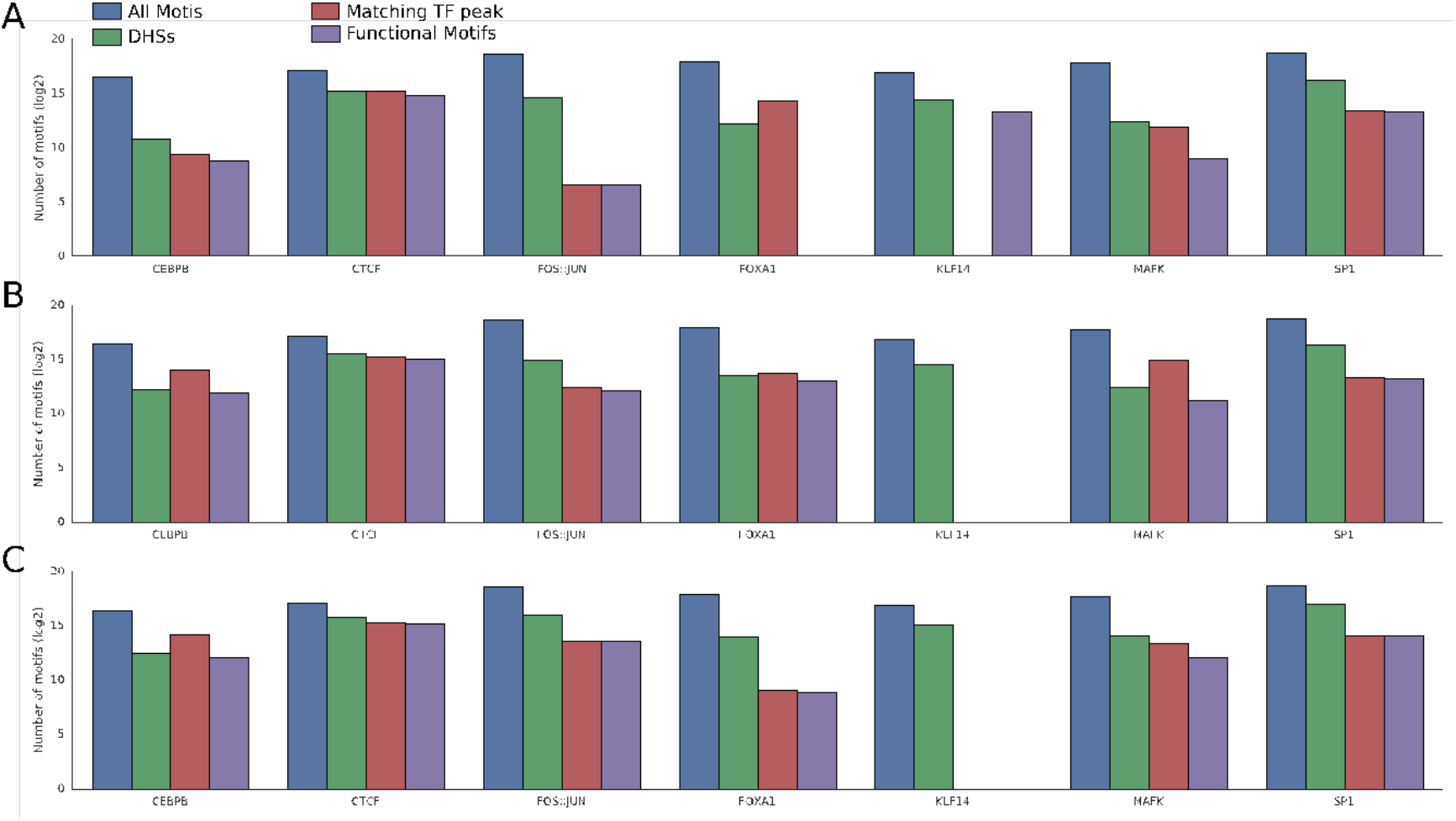
Motif-annotation overlaps (log2 scale) The columns represent number of sequences predicted motifs, motifs overlapping DHSs, motifs overlapping TF peaks and functional motifs respectively in (A) blood, (B) brain and (C) liver tissues.

**Supplementary Figure 2:**
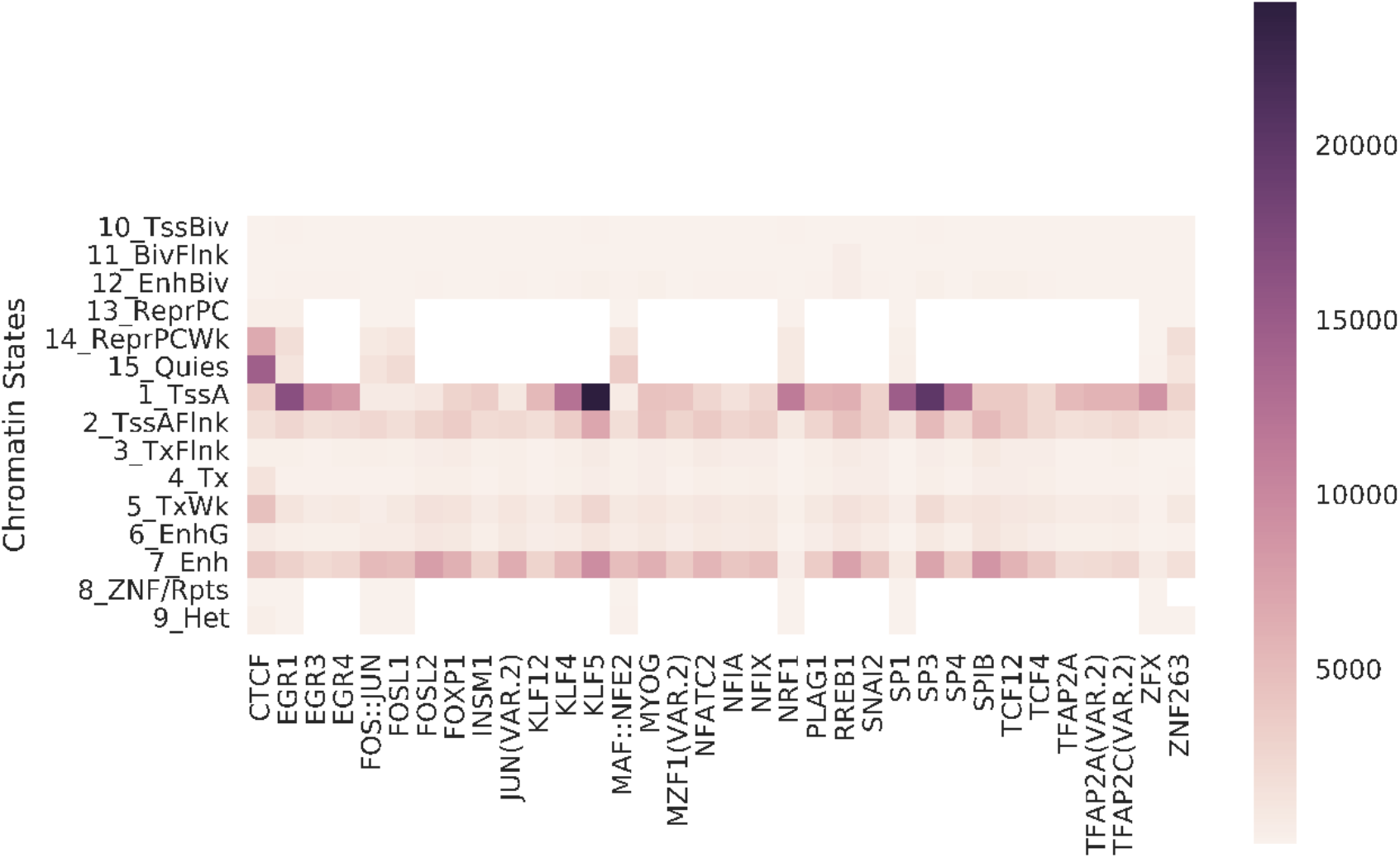
Enrichment of functional motifs in chromatin states in myeloid.

**Supplementary Figure 3:**
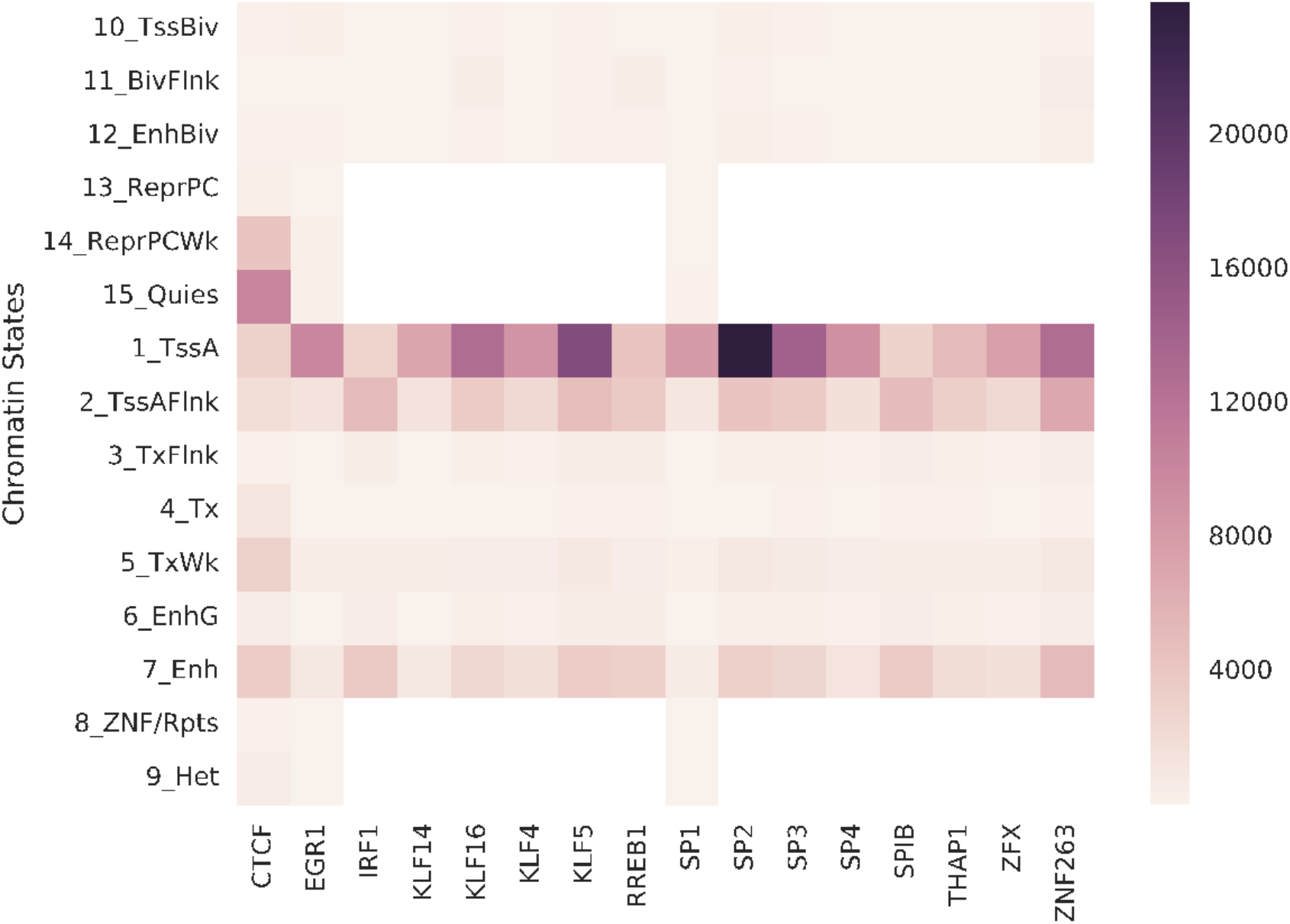
Enrichment of functional motifs in chromatin states in blood.

**Supplementary Figure 4:**
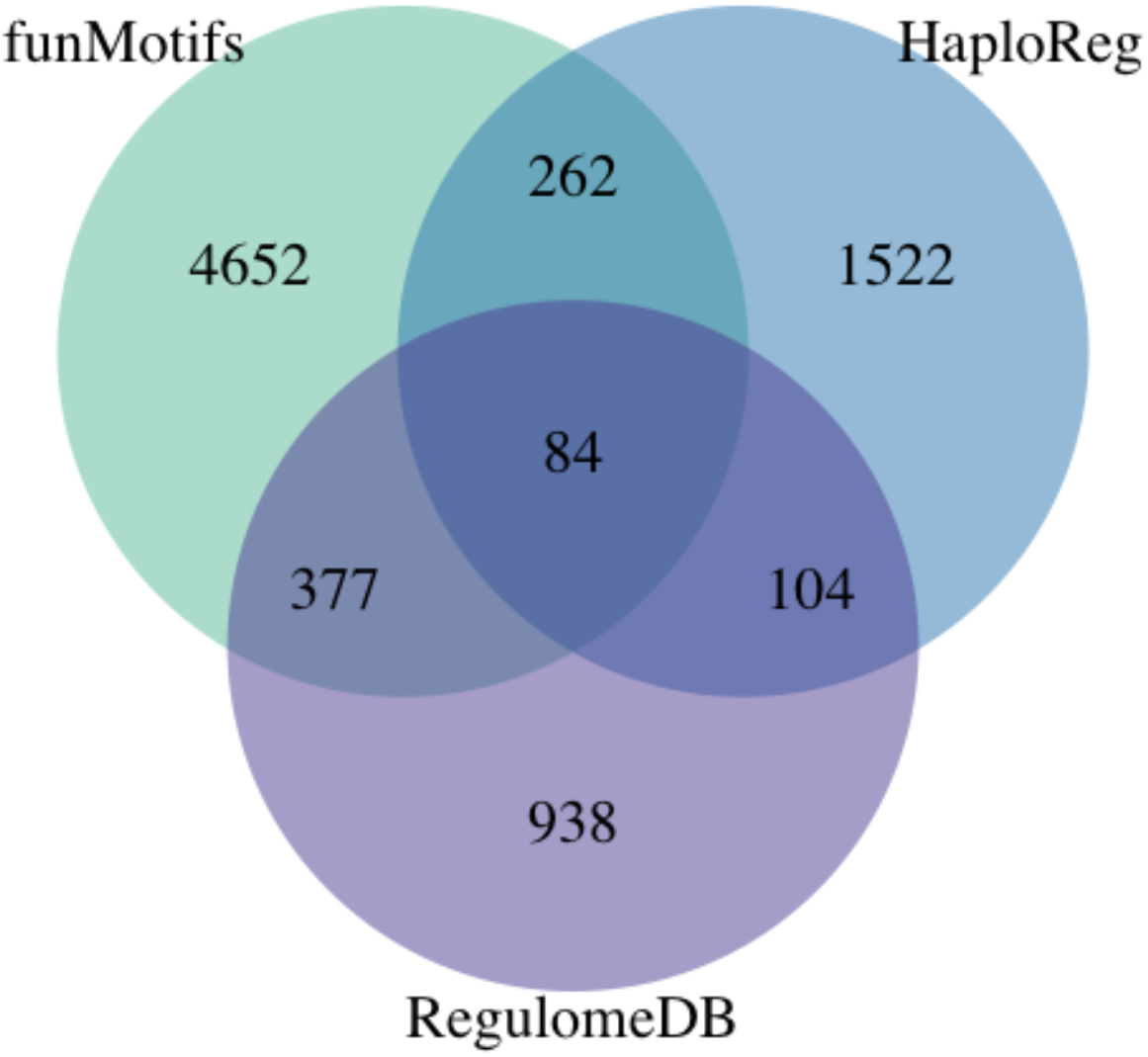
The comparison of candidate functional eQTL SNPs annotated by funMotifs, HaploReg and RegulomeDB.

**Supplementary Figure 5:**
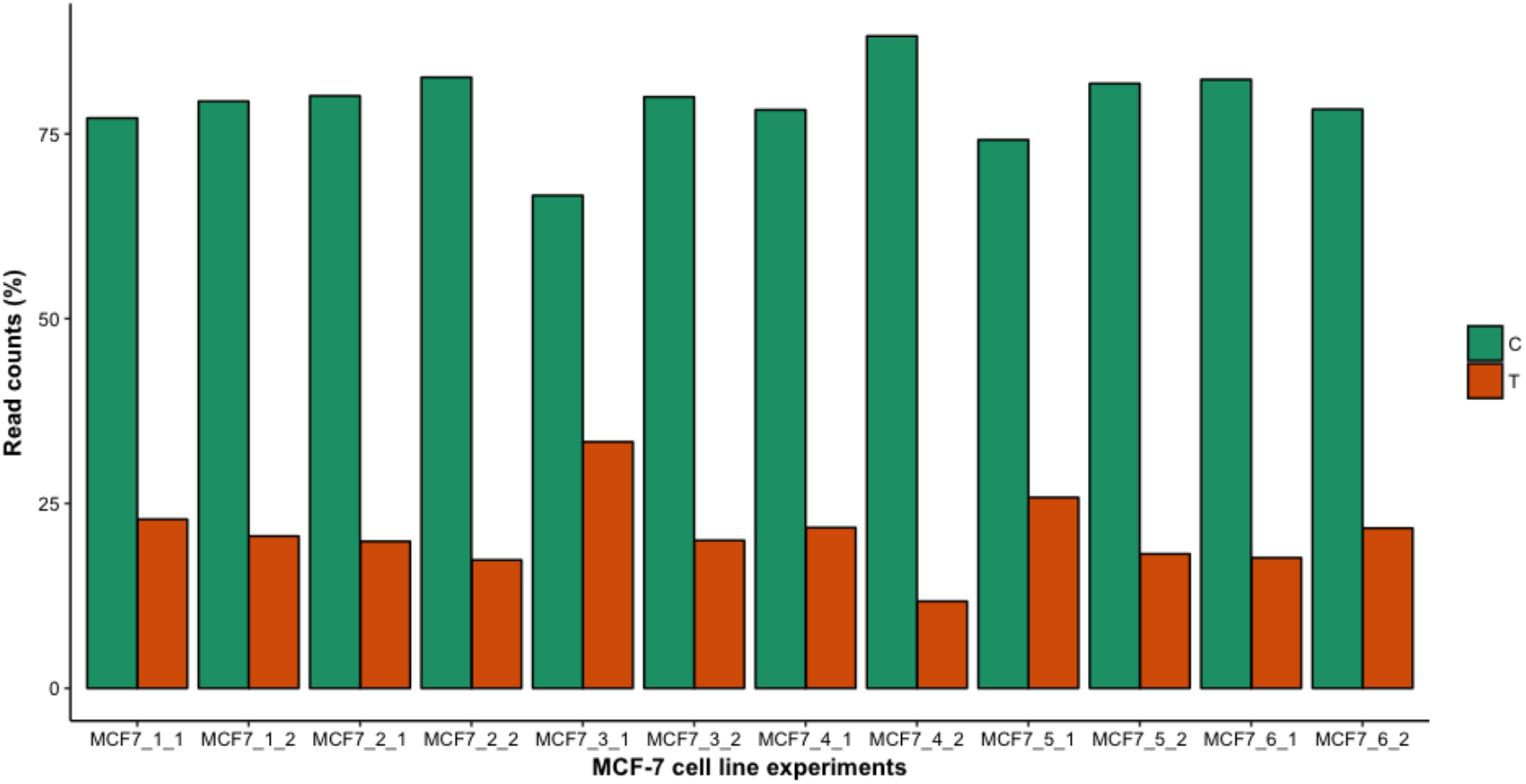
Read counts distribution for the candidate functional GWAS SNP rs8103622 for breast cancer cell line MCF-7 using ChIP-seq data (**Supplementary Table 13**).

**Supplementary Figure 6:**
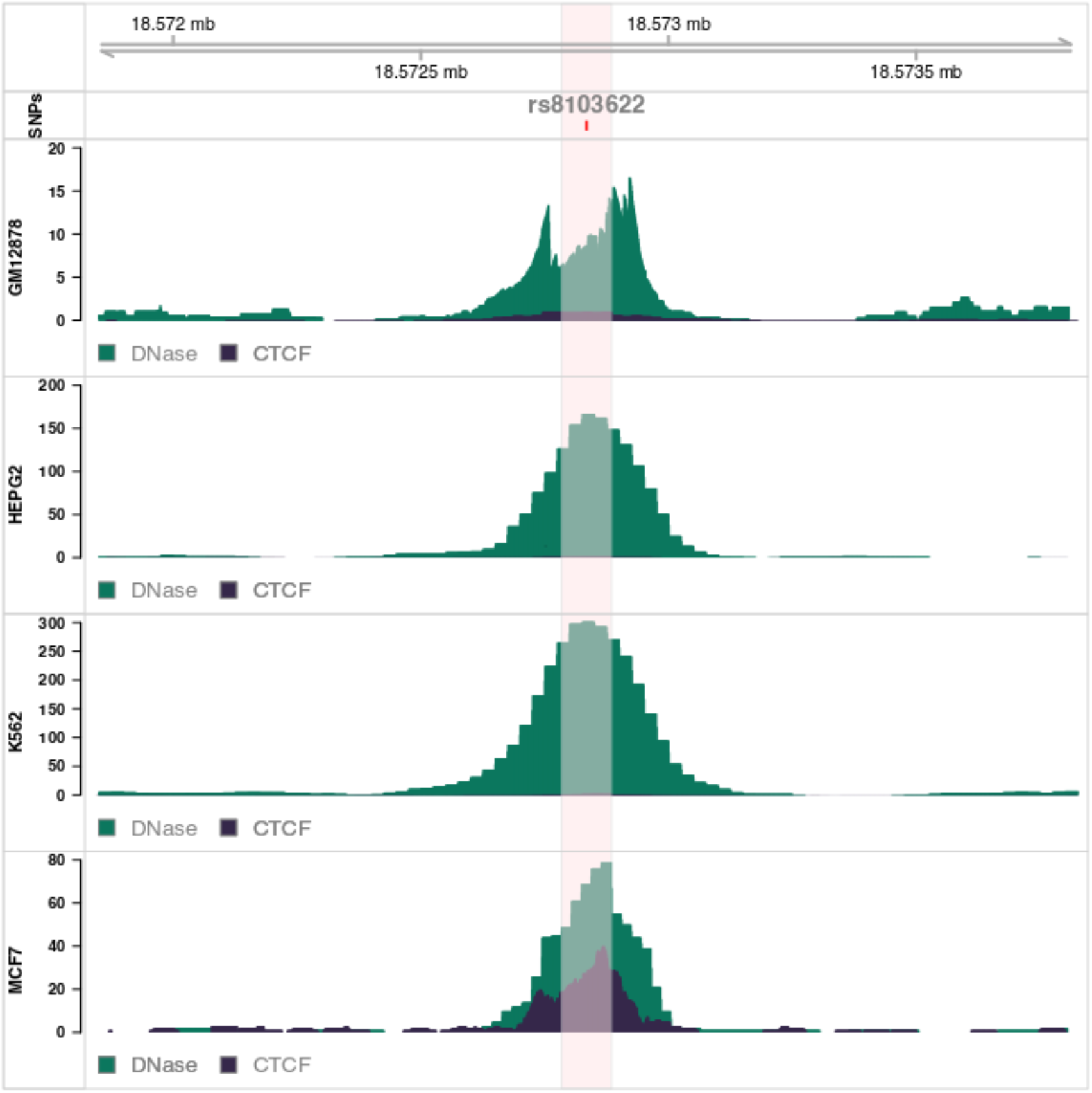
Presence of DHS and CTCF TF binding signals for the candidate functional GWAS SNP rs8103622 in four tissues: GM12878 – blood, HEPG2 – liver, K562 – myeloid and MCF7 – breast.

**Supplementary Figure 7:**
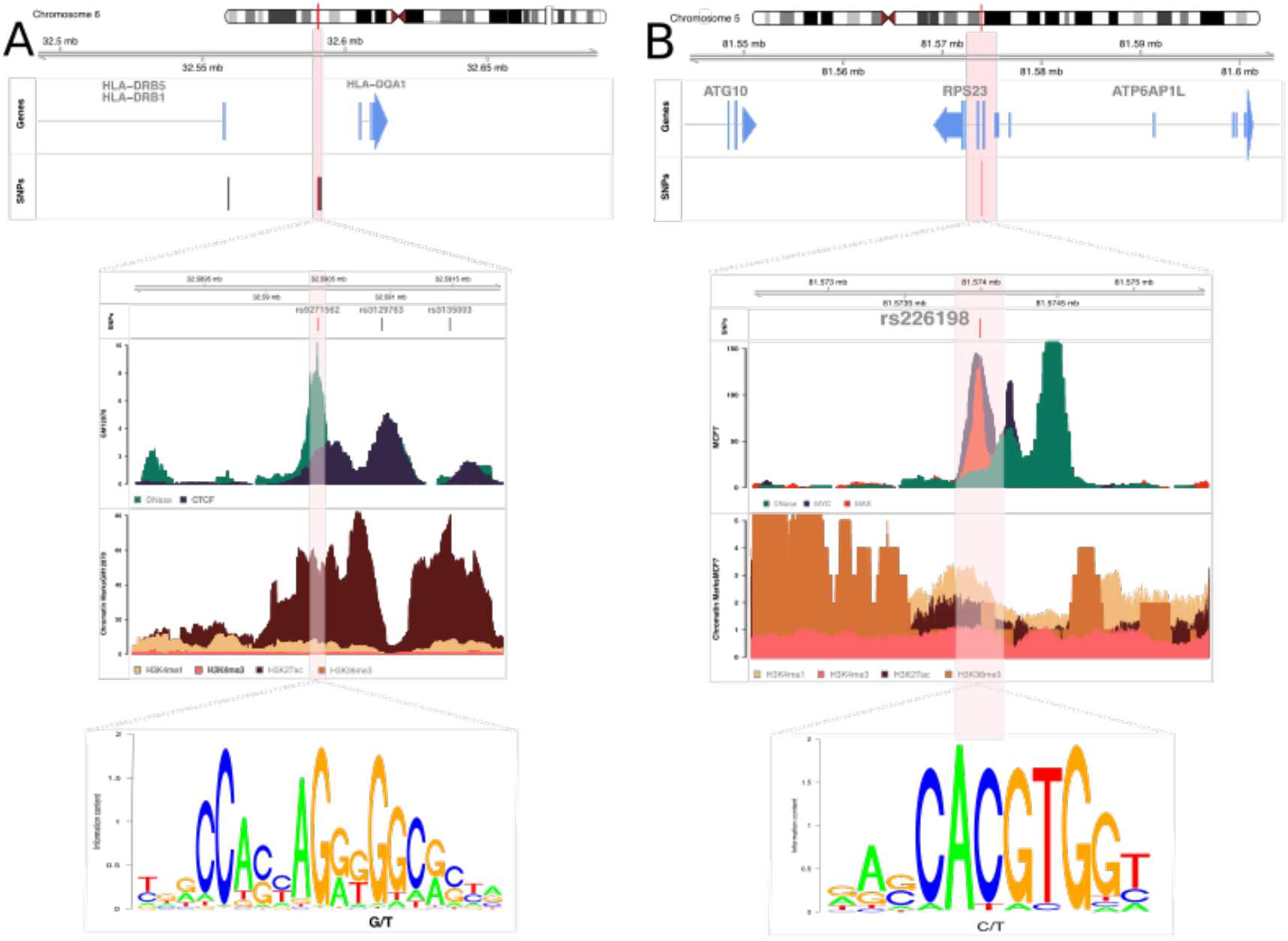
Overview of the candidate functional GWAS SNPs: a) rs9271562 in blood and b) rs226198 in breast. Genes box: blue genes are genes associated with the candidate SNP using eQTL SNPs. SNPs box: red line indicates position of the candidate SNP, black lines are GWAS SNPs in LD with the candidate SNP.

## Notes

http://bioinf.icm.uu.se/funmotifs

https://figshare.com/s/e6d3501d5a36617893b5

https://figshare.com/s/073c985fd131f78fb9c5

